# Programmable CRISPR interference for gene silencing using Cas13a in mosquitoes

**DOI:** 10.1101/846055

**Authors:** Aditi Kulkarni, Wanqin Yu, Alex Moon, Ashmita Pandey, Kathryn A. Hanley, Jiannong Xu

**Affiliations:** Department of Biology, New Mexico State University, PO Box 30001 MSC 3AF, Las Cruces NM, 88003, USA

**Keywords:** CRISPR-Cas13a, RNA interference, *Anopheles gambiae*, *Aedes aegypti*, gene silencing, CRISPRi

## Abstract

In the CRISPR-Cas systems, Cas13a is an RNA-guided RNA nuclease specifically targeting single strand RNA. We developed a Cas13a mediated CRISPR interference tool to target mRNA for gene silencing in mosquitoes. The machinery was tested in two mosquito species. A *Cas13a* expressing plasmid was delivered to mosquitoes by intrathoracic injection, and *Cas13a* transcripts were detectable at least10 days post-delivery. In *Anopheles gambiae*, *vitellogenin* gene was silenced by *Vg*-crRNA injection two hours post-blood meal, which was accompanied by a significant reduction in egg production. In *Aedes aegypti*, the α- and δ-subunits of *COPI* genes were silenced by a post-blood meal crRNA injection, which resulted in mortality and fragile midguts, reproducing a phenotype reported previously. Co-silencing genes simultaneously is achievable when a cocktail of target crRNAs is given. No detectable collateral cleavages of non-target transcripts were observed in the study. This study adds a programmable CRISPR tool to manipulate RNA in mosquitoes.

## Introduction

Characterization of mosquito life traits via functional genomics approaches can inform innovative control strategies through the identification of genes involved in development, host-seeking, blood feeding, digestion, fecundity, immunity, xenobiotic metabolism and insecticide resistance. The last is perhaps the most immediately impactful, as, with increasing insecticide resistance, the array of options for vector control is shrinking and in dire need of replenishment. RNA interference (RNAi) based approaches have been widely used to identify genes that are relevant to vector competence, and RNAi-based effectors for mosquito control have been developed ^1–5^. As an alternative or complement to RNAi-based tools, CRISPR-Cas9 based genome editing tools have been developed in the mosquito research field ^6–8^.

CRISPR-Cas systems are adaptive immune mechanisms used by prokaryotes to defend against invading DNA and RNA ^9–12^. Cas9 is a RNA-guided DNA nuclease and once assembled with a CRISPR guide RNA (sgRNA), Cas9 is able to cleave target DNA in a highly specific fashion, and DNA-targeting Cas9 has been harnessed for genome editing ^13, 14^. More recently, CRISPR-Cas9 based genome editing tools have been developed in the mosquito research field ^6–8^. Furthermore, catalytically inactive Cas9 (dCas9) was adapted for manipulation of gene expression. The dCas9 can be fused with a gene repressor or transcription activator. Guided by CRISPR RNA, such dCas9 proteins are able to bind target promoter or exonic DNA sequence without cleavage, and either repress transcription (CRISPR interference, CRISPRi) or activate transcription of target genes (CRISPR activation, CRISPRa) ^15–17^. Lately, Cas13 RNA nucleases ^10, 18, 19^, the new members in the CRISPR nuclease family, have been repurposed to specifically target endogenous RNAs as well as viral RNAs ^11, 20–23^. Most Cas13 proteins are single “effector” proteins with two Higher Eukaryotes and Prokaryotes Nucleotide-binding (HEPN) domains ^10, 24^. Once loaded with a target-specific crRNA, a Cas13 protein will locate target RNA and execute nuclease activity to degrade the target. Unlike Cas9, no Protospacer Adjacent Motif (PAM) sequences are required for Cas13 to function. Although a Protospacer Flanking Site (PFS), A, U, C, may be present for *Psp*Cas13b activity ^25^, no PFS is needed for *Lwa*Cas13a ^11^. The Cas13s tested for human RNA knockdown thus far have demonstrated high specificity and exhibited negligible, and significantly fewer, off-target effects than matched RNAi short hairpin RNAs (shRNA) using to trigger RNAi ^11, 25^. In bacteria, Cas13 HEPN-nuclease is able to cleave not only the target-RNA *in cis* but also other non-target RNA present *in trans* ^10^. Interestingly, no collateral effect was observed in three studies of CRISPR-Cas13 using human or plant cell lines ^11, 25, 26^, but Cas13a associated collateral RNA cleavage was reported in human glioma cancer cells ^27^. Notably, Cas13 can effectively silence several transcripts in parallel ^11, 20, 26, 28^. Taken together, CRISPR-Cas13 systems have become a new, exciting engine for CRISPRi ^29^.

In this study, we engineered a construct to express Cas13a from *Leptotrichia wadei* in mosquitoes and demonstrated its efficacy in silencing genes in two mosquito species, *Anopheles gambiae* and *Aedes aegypti*, that are major vectors of malaria and mosquito-borne viruses, respectively.

## Materials and Methods

### Plasmid construction

The Cas13a from *L. wadei* belongs to the class 2 type VI RNA-guided RNA nucleases ^11^. It’s RNA targeting effect has been demonstrated in human and plant cells^11, 20, 27, 30^. Plasmid pAc-sgRNA-Cas9 was used as a template to engineer construct pAc-Cas13a (**Fig. 1**). Plasmid pAc-sgRNA-Cas9 was a gift from Ji-Long Liu (Addgene plasmid # 49330). The codons of *LwaCas13a* gene were optimized to *Drosophila* preference. The codon optimized Cas13a sequence was synthesized and cloned at GenScript (https://www.genscript.com). The Cas9 coding sequence in plasmid pAc-sgRNA-Cas9 as replaced by the *Cas13a* coding sequence to create the pAc-Cas13a construct. The Cas13a sequence was confirmed by sequencing. The *Cas13a* was under the control of *Drosophila* actin (*Ac5*) promoter for constitutive *Cas13a* expression. The ampicillin resistance gene (*AmpR*) allows selection of positive clones (**Fig.1A**). The plasmid was replicated in *E.coli*, and extracted using a QIAprep Spin Miniprep kit (Cat No.27104, Qiagen). The sequence of plasmid pAc-LwaCas13a was deposited in NCBI under accession number (in the process, to be provided).

**Fig. 1.**
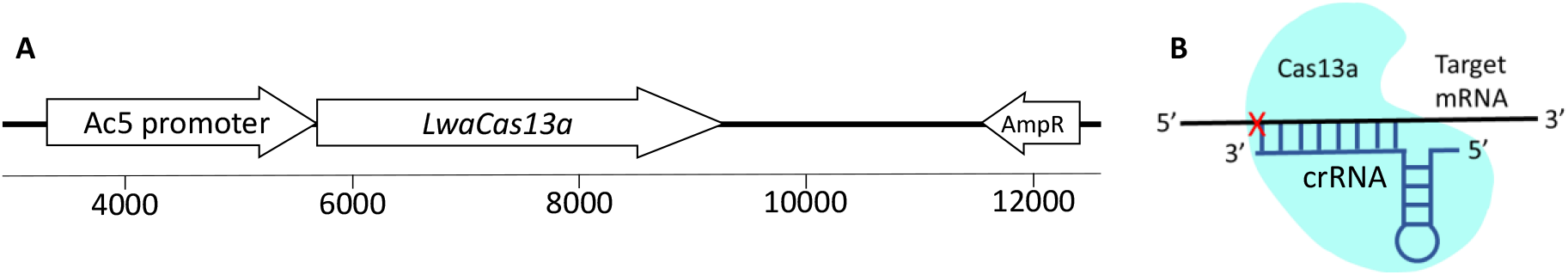
Cas13a construct and function. (A) Map of pAc-*Lwa*Cas13a (partial). (B) Mechanism of Cas13a/crRNA mediated target RNA cleavage.

### Synthesis of crRNA

A crRNA consists of a 36-nt direct repeat (DR) sequence and 28-nt target RNA specific sequence (N_28_), and the sequence of N_28_ is complementary to the target RNA sequence (**Fig.1 B**). The T7 promoter sequence (AGTTAATACGACTCACTATAGG) was added to the 5’ end of the DR sequence (GATTTAGACTACCCCAAAAACGAAGGGGACTAAAAC) to enable crRNA synthesis using T7 RNA polymerase *in vitro*. The target specific sequences (N_28_) used in this study are shown in **Table 1**. The selection of target (N_28_) is straightforward as *Lwa*Cas13a does not require PFS for activity ^11^. Please note, the target sequence for *COPI-α* was accidently designed as a-27nt sequence. For each *Vg*, *COPI-α* and *COPI-δ*, one target crRNA was used silencing, and for *Caspar* and *Cactus*, two target crRNAs were used. Template DNA duplexes of the crRNAs (T7-DR-N_28_) were synthesized at IDT Inc. (https://www.idtdna.com). The crRNAs were synthesized using T7-RNA polymerase (RPOLT7-RO ROCHE, Sigma-Aldrich). The crRNA synthesis reactions were set up in 40 μl containing template DNA duplex (1μg), 1 mM each of nucleotides ATP, GTP, CTP and UTP, 10X reaction buffer, T7 RNA polymerase 40U, and RNase inhibitor 20U. The reactions were incubated overnight at 37°C and stopped by heating the mixture at 65°C for 5 minutes. The crRNAs were treated with Turbo DNase I Kit (AM1907, ThermoFisher) to remove template DNA. The crRNA yield was quantified using a NanoDrop and stored at −20°C until use. Control crRNA (ctr crRNA) consisted of a randomly scrambled N_28_ nucleotide sequence, which had no homologous hit in the genomes of *An. gambiae* and *Ae. aegypti*.

**Table 1.**
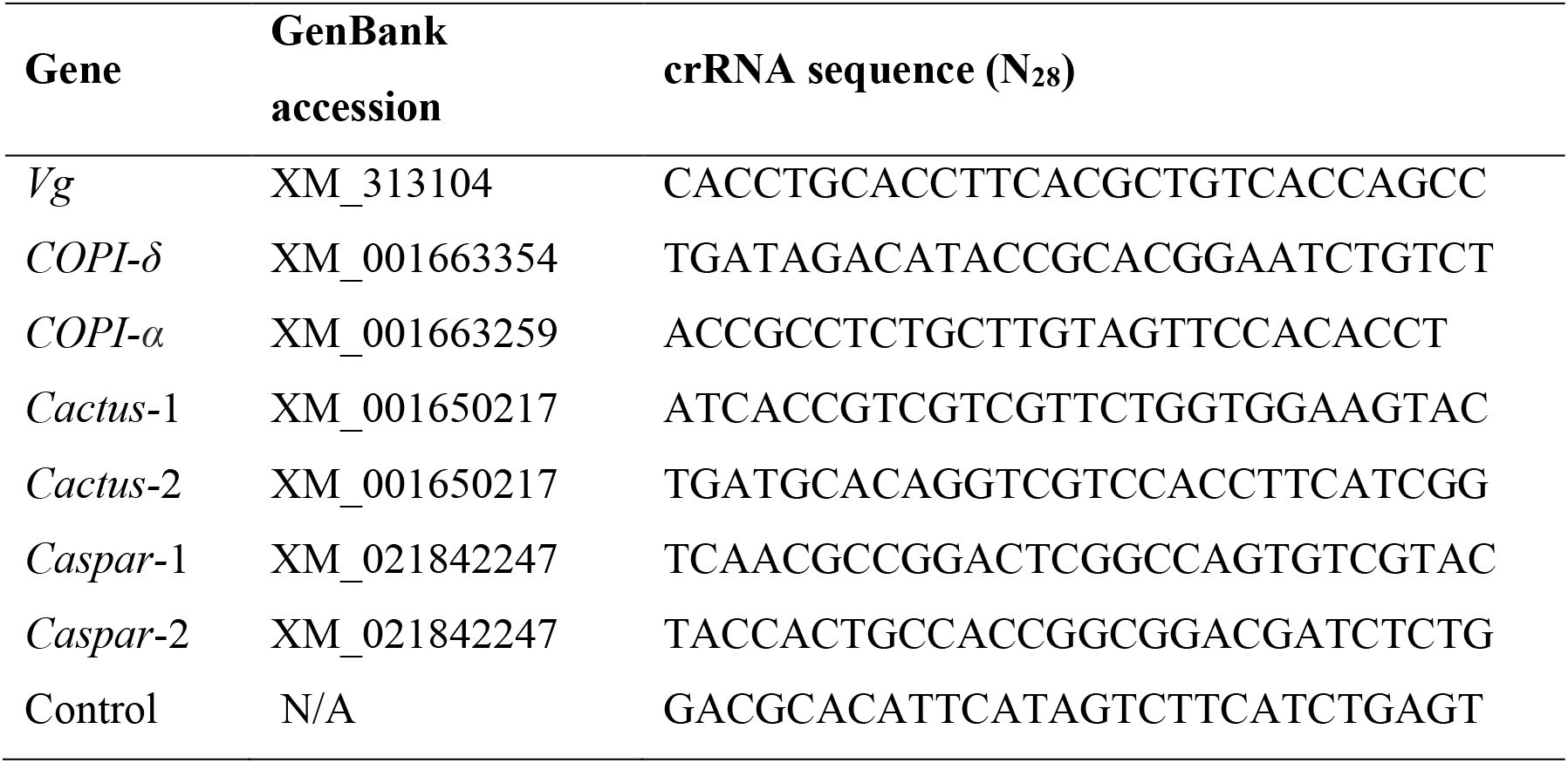
The crRNA (N_28_) sequences.

### Construct and crRNA Delivery

*An. gambiae* G3 strain and *Ae. aegypti* Puerto Rico strain were obtained from MR4 BEI and maintained using rearing conditions described previously ^31, 32^. The pAc-Cas13a construct (0.5μg/μl) was delivered into one-day old adult female mosquitoes by intrathoracic injection. To aid construct delivery into cells, the plasmid was mixed with a transfecting agent FuGENE HD (E2311, Promega) at concentration of 1.6μl of FuGENE reagent with 10μg construct DNA in 20 μl. The final concentration of construct DNA was 0.5 μg/μl in the mixture. Approximately, each *An. gambiae* mosquito received 100 nl mixture, and each *Ae. aegypti* mosquito received 150 nl mixture. Gene specific crRNAs were either delivered with the construct or separately at a later time point. For blood inducible genes, *Vg* and *COPI*, female mosquitoes were given a blood meal three days post construct injection. Corresponding crRNAs (0.5μg/μl, prepared in FuGENE as described above) were intrathoracically injected into mosquito hemocoel at two hours post blood meal.

### RNA isolation, cDNA synthesis and PCR

Total RNA from whole mosquitoes was isolated using Trizol (Invitrogen) following the manufacturer’s instruction. The RNA was treated with Turbo DNase I Kit to remove genomic DNA contamination, and then 1μg RNA was converted to cDNA using Protoscript II RT (M0368S, New England Biolabs) following the manufacturer’s instruction. The PCR assays were performed using 1μl 1:5 diluted cDNA as template, 0.2 μM primers (primer sequences are presented in Table S1) and 2 × PCR Master mix (M0482S, NEB), with the following cycling parameters: 35 cycles of denaturing at 95°C for 15 seconds, annealing at a temperature optimal for the amplicon (Table S1) for 15 seconds, and extension at 68°C for 20 seconds with an extra 5 min in the last cycle for final extension.

### Statistical analysis

In the *Vg* gene knockdown experiment, eggs were dissected from ovaries at day 3 post blood meal. The egg counts were compared between the *Vg* crRNA and control crRNA cohorts. The non-parametric Mann-Whitney test was used for statistical comparison of the egg numbers. In the *COPI* gene knockdown experiment, a survival curve was plotted using GraphPad Prism, and a Mantel-Cox analysis was performed to compare the survival between the *COPI* crRNA and control crRNA cohorts.

## Results

### *Cas13a* expression in mosquitoes post intrathoracic delivery

A construct was engineered to express *Cas13a* gene by modifying the plasmid pAc-sgRNA-Cas9, which was successfully used to transfect *Drosophila* cells for targeted genetic mutagenesis previously by Bassett et al. (2014) ^33^. As shown in **Fig. 1**, Cas13a coding sequence is under the control of constitutive promoter Ac5. The plasmid was prepared with transfection reagent FuGENE HD and injected into thorax of one-day old mosquitoes. *Cas13a* transcription in whole mosquitoes was determined by RT-PCR. In *An. gambiae*, the RNA was sampled at day three post construct delivery, and *Cas13a* transcripts were detected (**Fig. 2A**). No *Cas13a* amplification was observed in the controls that did not receive the construct. In *Ae. aegypti*, the RNA samples were collected on day 2, 5, 7 and 10 post construct delivery, and *Cas13a* transcripts were detected in all of these time points (**Fig. 2B**).

**Fig. 2.**
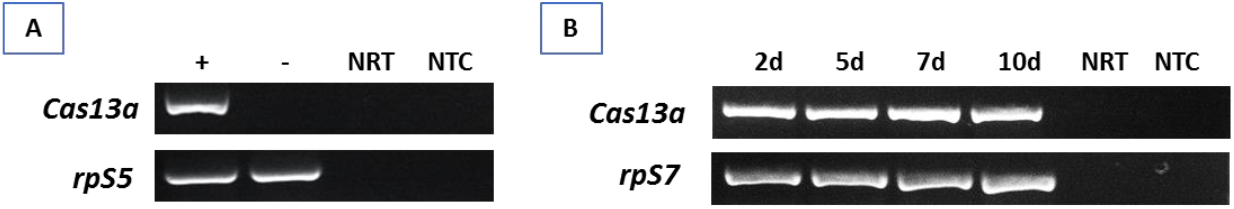
Detection of *Cas13a* transcript in pAC-Cas13 injected mosquitoes. (A) RT-PCR of *Cas13a* in *An. gambiae* mosquitoes that received the construct (+) vs. control (−) at day 3 post plasmid delivery. NRT: No RT control; NTC: no template control. The *rpS5* was used as a loading control. (B) Expression pattern of *Cas13a* transcript over a period of 10 days in *Ae. aegypti*, the *rpS7* was used as a loading control.

### Cas13a mediated *Vitellogenin* gene silencing in *An. gambiae*

In mosquitoes, yolk protein precursor vitellogenins (Vg) are required for the vitellogenic stage in oogenesis after blood feeding ^34^. To test Cas13a/crRNA mediated *Vg* silencing, the Cas13a construct was delivered to one-day old *An. gambiae* (N = 120). Three days later, the mosquitoes were given a blood meal to induce *Vg* expression (N = 96). To activate *Vg* knockdown, *Vg* crRNA (N = 41) or control crRNA (N = 40) were injected into the blood engorged females at 2 hr post feeding. The *Vg* RT-PCR was used for verification of *Vg* knockdown. As shown in **Fig. 3**, the abundance of *Vg* transcript was reduced in females that received *Vg*-crRNA as compared to females that received control-crRNA. As expected, successful *Vg* knockdown resulted in reduction in egg production (**Fig. 3**). The *Vg*-crRNA treated mosquitoes produced on average 39 ± 25 (mean ± SD) eggs/female (N = 33), while control mosquitoes produced 64 ± 23 eggs/female (N = 32). The difference was statistically significant (Mann-Whitney test, P<0.001). A second experimental replicate generated data with similar pattern showing significant reduction in egg numbers (Mann-Whitney test, P<0.01, **Fig. S1**). The Vg proteins are produced by fat body and secreted into hemolymph, and subsequently deposited in developing oocytes via receptor-mediated endocytosis. The results suggest that the Cas13 construct was successfully delivered into fat body cells, where the *Cas13a* was transcribed and translated into functional Cas13a proteins. Post blood feeding, *Vg*-crRNAs were delivered into cells, where they were assembled with Cas13a to form a target specific RNA nuclease complex, which cleaved the *Vg* mRNA effectively.

**Fig. 3.**
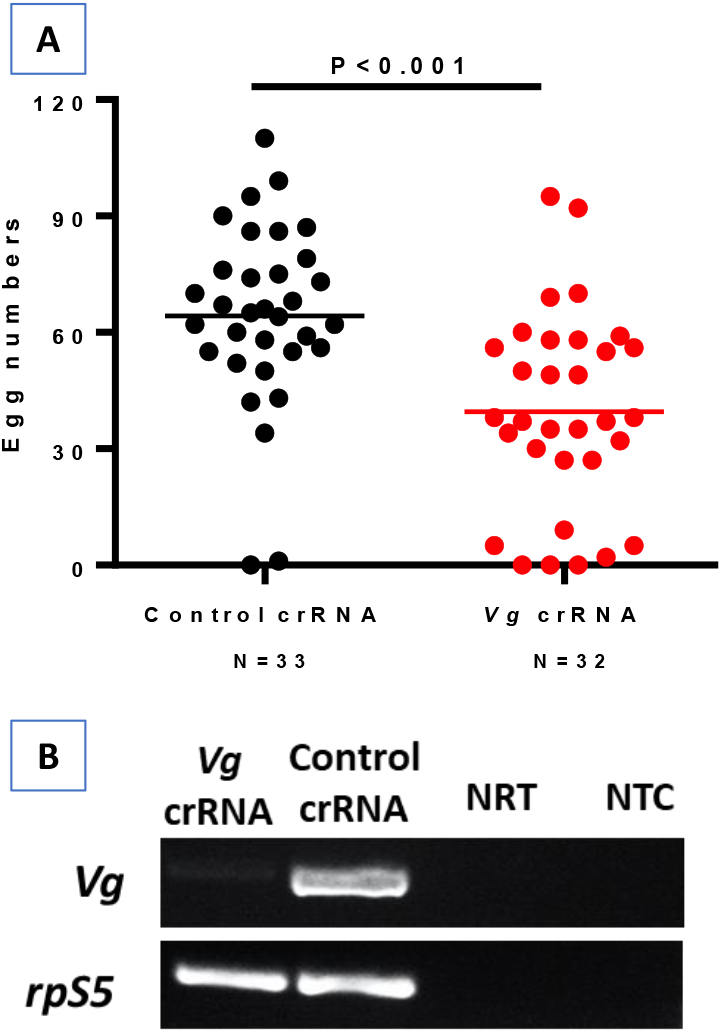
*An. gambiae Vg* knockdown reduced egg production. (A) Mosquitoes treated with Cas13a/*Vg-crRNA* had lower egg counts as compared to the cohort treated with Cas13a/ctr-crRNA (P<0.001). (B) Reduction of *Vg* transcripts was confirmed by RT-PCR. NRT: No RT control; NTC: no template control. The *rpS5* was used as a loading control.

### Cas13a mediated *COPI* gene silencing in *Aedes aegypti*

The coatomer complex I (COPI) proteins are involved in blood digestion in *Ae. aegypti* ^35^. The COPI complex consists of α, β, β’, γ, δ, ε and ζ subunits, each encoded by separate genes. The production of COPI proteins is blood inducible between 18-36 hr post blood meal in fat body and 24-48 hr post blood meal in ovaries ^35^. As shown by other investigators, dsRNA-mediated knockdown of the genes encoding all but the ε subunit led to blood meal-induced mortality ^35^. Therefore, we targeted the *COPI* genes to test the gene silencing efficacy of the Cas13a machinery in *Ae. aegypti.* The Cas13a construct was injected into one-day old mosquitoes (N = 90). At day 3 post construct delivery, the mosquitoes were given a blood meal. The engorged mosquitoes (N = 75) were split into two cohorts, one was injected with the crRNAs specific for α and δ*COPI* genes at 2 hr hours post blood meal (N = 27). The other cohort (N = 29) was injected with the control crRNA. A subset of mosquitoes (N = 5) in each cohort was sampled for knockdown validation by RT-PCR at 20 hr post blood meal. Reduced abundance of the *COPI* transcripts was observed in the knockdown group (**Fig. 4**). The survival curves over 9 days post blood meal revealed a significantly lower survival of the *COPI* knockdown cohort (N = 22) than the control cohort (N = 24) (**Fig. 4**; Mantel-Cox test, P<0.001). It has previously been shown that *COPI* knockdown makes the mosquito midgut fragile ^35^. Consistently, we observed that the midguts in the *COPI* knockdown mosquitoes were apt to break and leak during dissection, while the midguts of control mosquitoes were in good shape with intact blood bolus (**Fig. 4**). A second, replicate experiment also showed a significant reduction in the survival of the *COPI* knockdown cohort (Mantel-Cox test, P<0.01, **Fig. S2**).

**Fig. 4.**
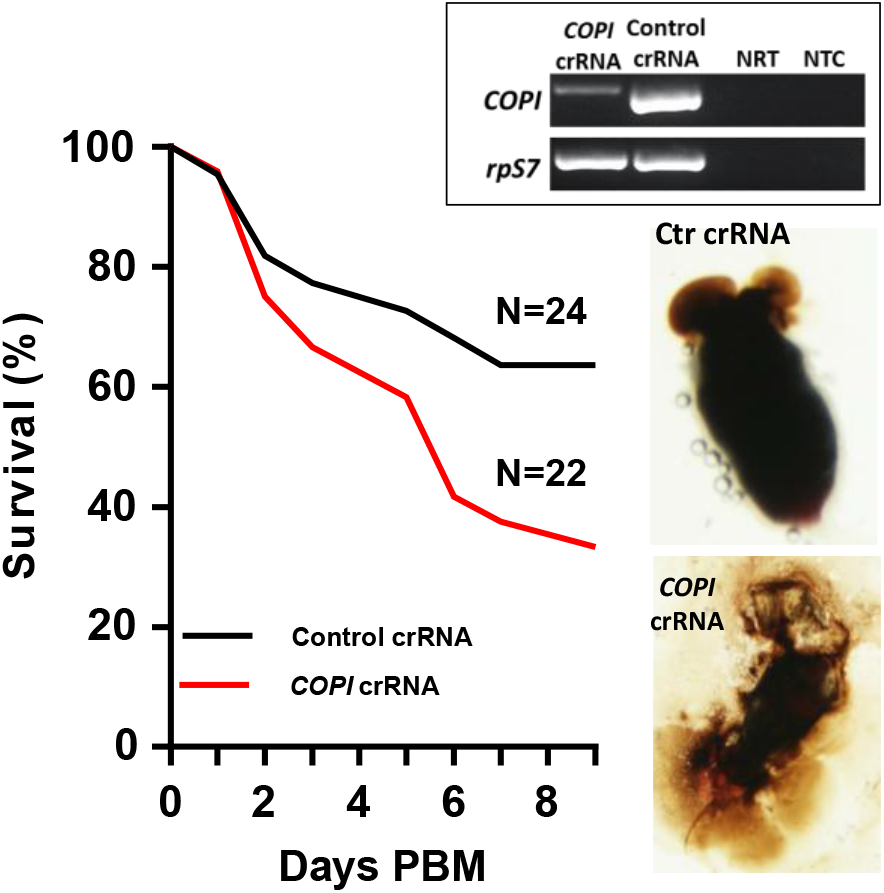
Phenotypes of *Ae. aegypti COPI* knockdown. Cas13a/*COPI*-crRNA resulted in mortality post blood meal (PBM) and fragile midgut. Reduction of *COPI* transcripts was confirmed by RT-PCR (insert panel). NRT: No RT control; NTC: no template control. The *rpS7* was used as a loading control.

### Cas13a mediated double gene knockdown

To determine the potential for silencing multiple genes, a cocktail of the Cas13a construct and crRNAs against the genes *Cactus* and *Caspar* or *Cactus* and *COPI* was delivered into *Ae. aegypti.* As expected, the target gene transcripts were reduced effectively by the respective treatment (**Fig. 5**).

**Fig 5.**
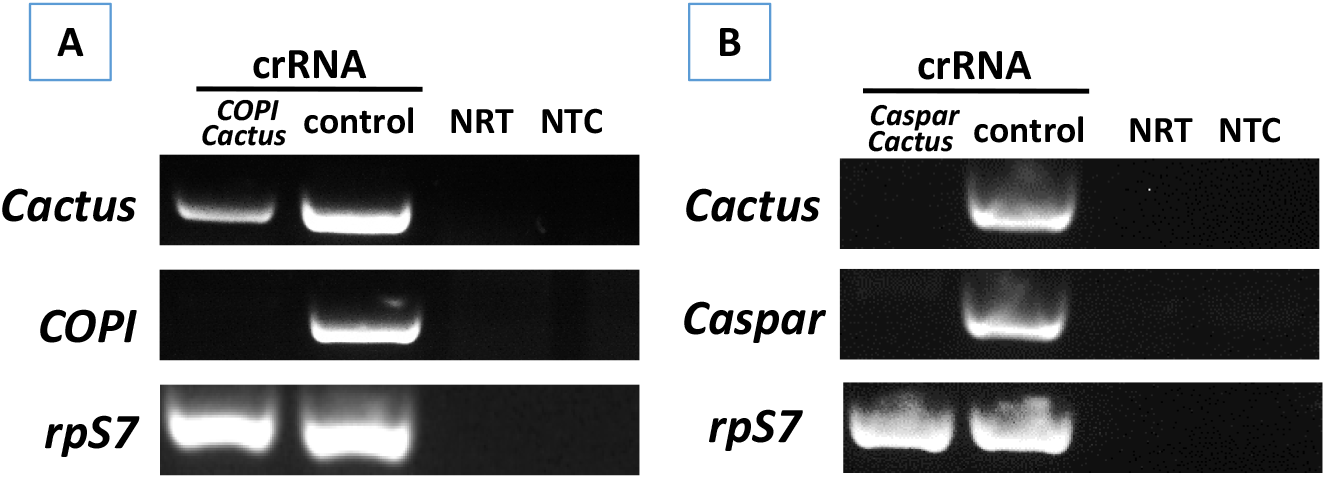
**RT-PCR verification of double gene knockdown** in *Ae. aegypti* that received a cocktail of Cas13a construct with (A) *Cactus*-and *Caspar*- or (B) *Cactus*- and *COPI*-crRNAs. NRT: No RT control; NTC: no template control. The *rpS7* was used as a loading control.

### Absence of detectable collateral cleavage of non-target RNA

In bacteria, activated Cas13a displays cleavage activity of non-target RNA^10^. To determine if Cas13a has such collateral activity in mosquitoes, we examined the abundance of arbitrarily selected non-target transcripts in Cas13a/*Vg* crRNA treated *An. gambiae* as well as Cas13a/*COPI-Cactus* crRNA treated *Ae. aegypti.* As shown in **Fig. 6**, in *An. gambiae*, Cas13a/*Vg* crRNA treatment resulted in *Vg* knockdown, but non-target transcripts *rpS5*, *rpS7* and *GAPDH* (glyceraldehyde 3-phosphate dehydrogenase) were not affected by the activated Cas13a. Likewise, in *Ae. aegypti, COPI/Cactus* crRNA activated Cas13a to co-silence *COPI* and *Cactus*, but did not affect non-target transcripts *Caspar, rpS7* and *rpS17* (**Fig. 6**). The data indicate that in mosquitoes the Cas13a may not execute collateral cleavage activity on non-target RNA.

**Fig 6.**
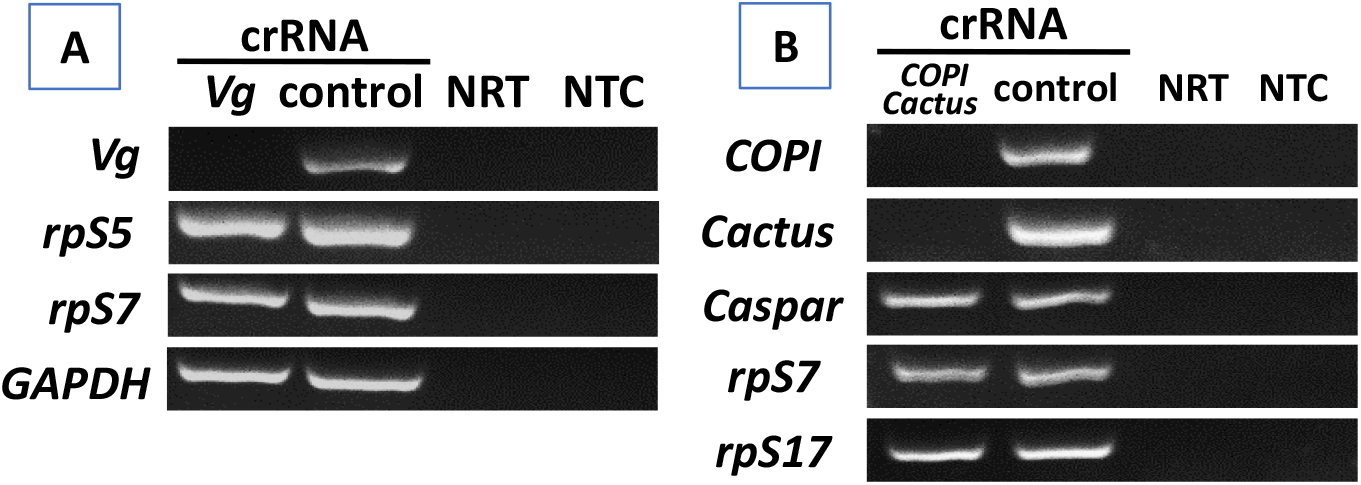
RT-PCR of non-target transcripts. (A) In *Vg* silenced *An. gambiae*, the abundance of non-target transcripts *rpS5, rpS7*, and *GAPDH* was not affected by Cas13a/*Vg* crRNA. (B) In *COPI* and *Cactus* co-silenced *Ae. aegypti*, the abundance of non-target transcripts *Caspar, rpS7*, and *rpS17* was not affected by Cas13a/*COPI-Cactus* crRNA. NRT: No RT control; NTC: no template control.

## Discussion

In this proof of concept study, we demonstrate the effectiveness of CRISPRi mediated by CRISPR-Cas13a/crRNA machinery. The *LwaCas13a* was derived from *L. wadei*, and under control of *Actin* promoter (**Fig.1**). The construct was delivered into the hemocoel of adult mosquitoes by intrathoracic injection, and *Cas13a* was transcribed constitutively (**Fig. 2**). Likely, the construct enters into the nucleus where the *Cas13a* gene is transcribed, and then the mRNA comes into the cytoplasm where the protein is translated. The construct remained active to transcribe *Cas13a* for at least 10 days post-delivery in *Ae. aegypti* (**Fig. 2**), which makes it temporally flexible to administer crRNAs targeting various genes that are expressed at different time points during a mosquito’s life span. Target-specific crRNAs can be delivered either with the construct together or after the construct delivery as appropriate to the experimental design. The system can silence highly abundant transcripts, as demonstrated by targeting *Vg* transcripts in *An. gambiae* (**Fig. 3**) and *COPI* transcripts in *Ae. aegypti* (**Fig. 4**). In both cases, the target genes are induced by blood meal to a high transcriptional abundance. In addition, we have tested the system on the genes *Cactus* and *Caspar* in *Ae. aegypti* (**Fig. 5)**. Taken together, these data conclusively demonstrate that the Cas13a-CRISPRi machinery is functional in *An. gambiae* and *Ae. aegypti* mosquitoes we tested in this study. This tool may work well in other mosquito species. We have not tested knockdown effect of the Cas13a system on genes that are mainly expressed in midgut, salivary glands and ovaries, which needs further studies. It would be an appropriate approach to develop a transgenic line expressing *Cas13a* gene for such studies.

The Cas13a-CRISPRi has certain advantages over the dCas9-CRISPRi for repressing gene expression ^16, 36^. First, the dCas9-CRISPRi machinery acts at DNA level while Cas13a targets mRNA directly. RNA-guided binding of dCas9 to a specific promoter or coding sequence can block transcription. This mode of action is efficient in bacteria, but often is not very efficient in eukaryotic cells ^16^. The dCas9 fusion proteins with a repressive domain have been developed for transcriptional repression, such as dCas9-KRAB (Krüppel associated box) in mammalian cells ^15^, but it is challenging to develop a fusion dCas9 with universal applicability. In addition, target specific sgRNA selection may be limited by the PAM that is required for the Cas9 activity. On the contrary, no PFS is required for target RNA cleavage by *Lwa*Cas13a in eukaryotic cells ^23^. The mode of action of Cas13 is simple and programmable, and the efficacy has proven high in mammalian cells ^11, 25^ and in mosquitoes in the current study. RNAi mediated gene silencing is a very common practice in mosquito gene function studies. Recently, application versions have been developed for mosquito vector control ^4, 5^. In these application cases, dsRNA is used to trigger RNAi machinery. In dsRNA mediated RNAi, the effective siRNA sequences sometimes are difficult to predict, therefore, large dsRNA fragments are often used to increase chances to generate effective siRNA by Dicer. However, this strategy is accompanied with a higher chance to produce siRNA with off-target potentials ^37, 38^. In addition, efficacy of dsRNA mediated RNAi varies case by case, and an optimal outcome is often a result of an empirical process ^39^.

There is a concern about the potential of collateral cleavage with Cas13a, in which non-target RNA sequences can be cleaved by Cas13a in bacteria ^10, 11^. Interestingly, this promiscuous RNA degradation activity was not observed in several studies in mammalian and plant cells ^11, 25, 26^. These data have warranted its safety to be used as an effector to target against RNA viruses that infect humans ^23^. However, a Cas13a/crRNA associated collateral cleavage was recently shown in human U87 glioblastoma cells ^27^. In the study, exogenous gene *GFP* or *EGFRVIII* were overexpressed and targeted by Cas13a/crRNA. In this context, a partial degradation of ribosomal RNA profile was observed, and the abundance of non-target transcripts of *GAPDH, HOTHAIR* and *L3MTL1* was reduced as well ^27^. Furthermore, the RNA integrity was compared between the LN229 glioma cell line and HEK293T cells after treatment with the Cas13a/crRNA, the LN229 cells tended to be more sensitive to the collateral effect than the HEK293T cells ^27^. In our study, we examined arbitrarily selected non-target transcripts, three in *Vg* silenced *An. gambiae* and three in *COPI* and *Cactus* co-silenced *Ae. aegypti;* no detectable reduction of these non-target mRNAs was observed (**Fig. 6**). The data suggest that collateral effect of Cas13a may not be a concern in mosquitoes, although we cannot completely rule out the possibility of collateral cleavage. Additional studies with larger scale examination of non-target RNA would help confirm the absence of non-target cleavage by the Cas13/crRNA system in mosquitoes.

The Cas13a-CRISPRi system holds promise for robust and flexible programming to silence one or more genes simultaneously in mosquitoes, with potential applications in other arthropods.

## Acknowledgments

This research is supported by an Institutional Development Award (IDeA) from the National Institute of General Medical Sciences of the National Institutes of Health under grant number P20GM103451, the National Institutes of Health SC1AI112786 and the National Science Foundation [No. 1633330]. The content is solely the responsibility of the authors.

## Authorship Confirmation Statement

JX conceived the idea for the project and devised the study. JX, AK, KAH designed experiments, AK, WY, ASM, AP conducted experiments. AK, JX, KAH wrote manuscript. The authors confirm that all co-authors have reviewed and approved the manuscript. The authors affirm that the paper is original with unpublished findings, not under consideration by any other journals.

---

**Table S1.**
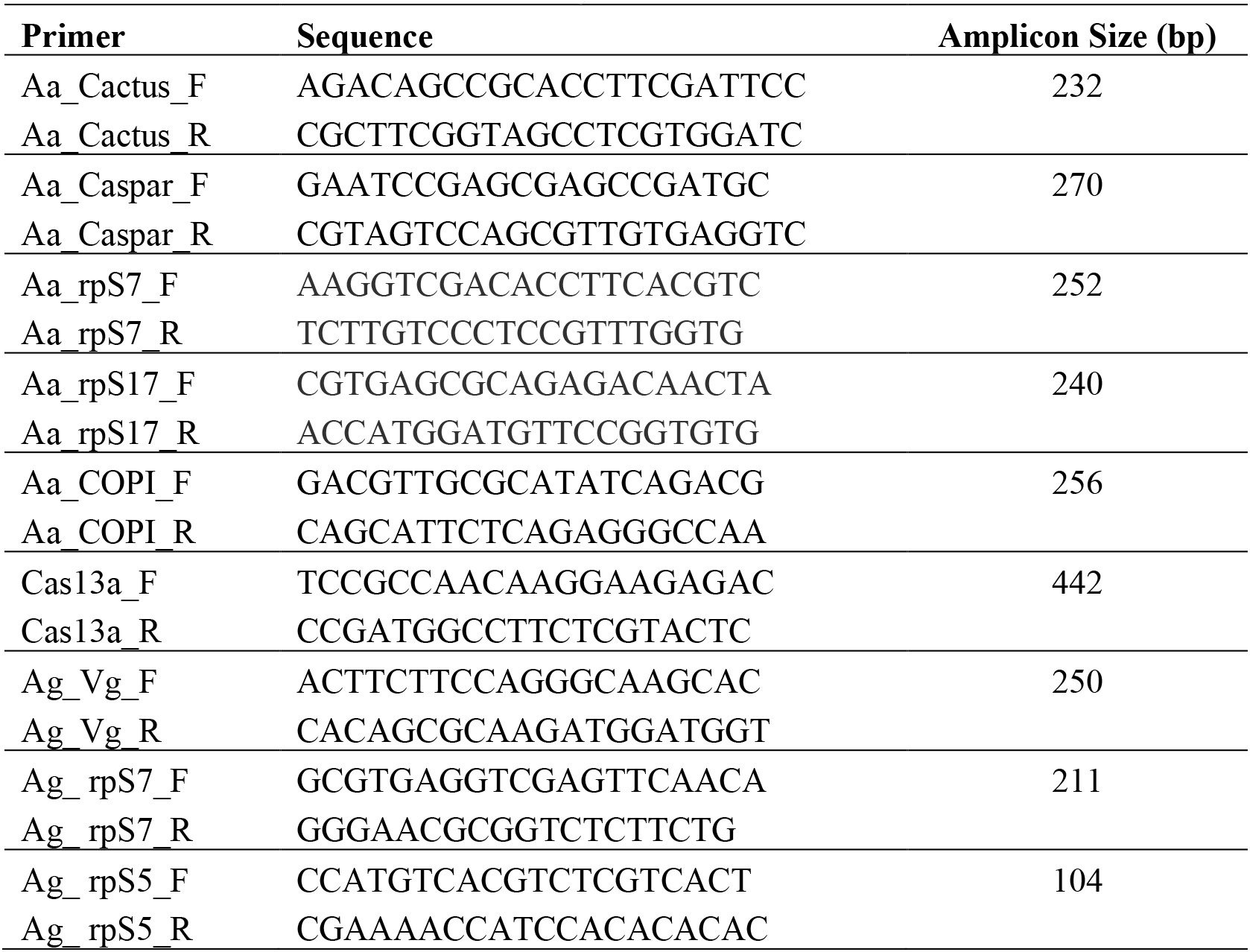
Primer sets used in the study.

**Figure S1.**
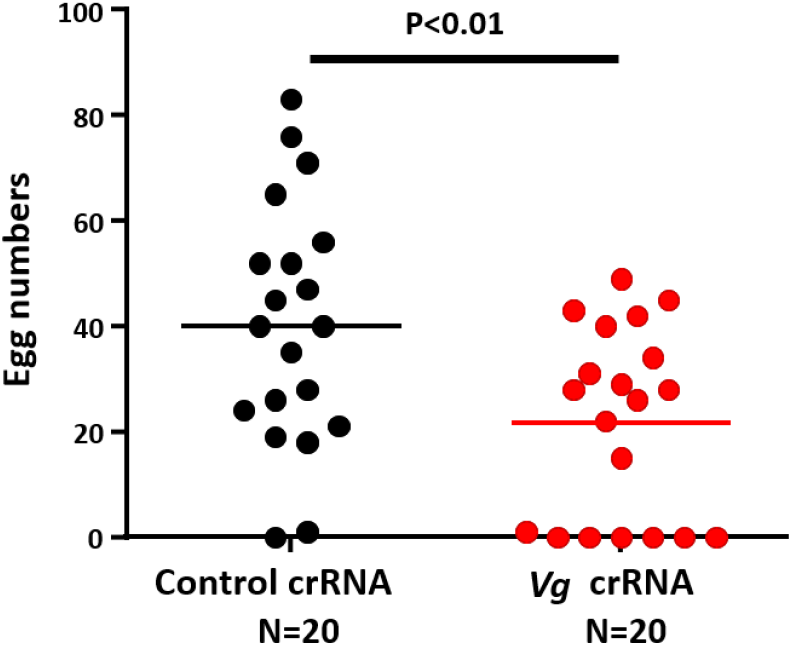
The Cas13a/*Vg* crRNA treatment significantly reduced egg production in *An. gambiae*. Mann-Whitney test, P<0.01.

**Figure S2.**
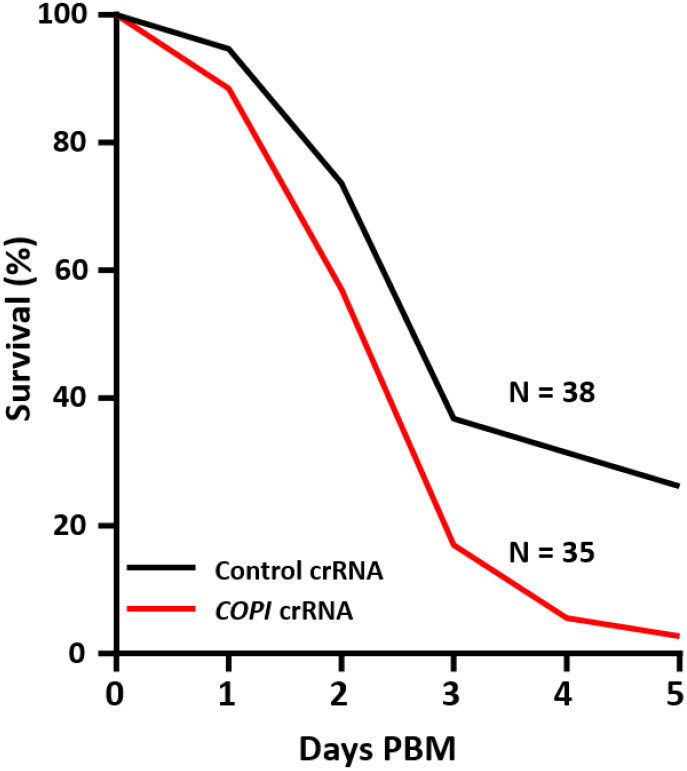
The Cas13a/*COPI*-crRNA treatment resulted in a significant higher mortality post a blood meal. Mantel Cox test, P<0.01.

