## Supplementary material for "Programmable CRISPR interference for gene silencing using Cas13a in mosquitoes": Table S1

| **Table S1. Primer sets used in the study** | | |
| --- | --- | --- |
| **Primer** | **Sequence** | **Amplicon Size (bp)** |
| Aa_Cactus_F | AGACAGCCGCACCTTCGATTCC | 232 |
| Aa_Cactus_R | CGCTTCGGTAGCCTCGTGGATC |  |
| Aa_Caspar_F | GAATCCGAGCGAGCCGATGC | 270 |
| Aa_Caspar_R | CGTAGTCCAGCGTTGTGAGGTC |  |
| Aa_rpS7_F | AAGGTCGACACCTTCACGTC | 252 |
| Aa_rpS7_R | TCTTGTCCCTCCGTTTGGTG |  |
| Aa_rpS17_F | CGTGAGCGCAGAGACAACTA | 240 |
| Aa_rpS17_R | ACCATGGATGTTCCGGTGTG |  |
| Aa_COPI_F | GACGTTGCGCATATCAGACG | 256 |
| Aa_COPI_R | CAGCATTCTCAGAGGGCCAA |  |
| Cas13a_F | TCCGCCAACAAGGAAGAGAC | 442 |
| Cas13a_R | CCGATGGCCTTCTCGTACTC |  |
| Ag_Vg_F | ACTTCTTCCAGGGCAAGCAC | 250 |
| Ag_Vg_R | CACAGCGCAAGATGGATGGT |  |
| Ag_ rpS7_F | GCGTGAGGTCGAGTTCAACA | 211 |
| Ag_ rpS7_R | GGGAACGCGGTCTCTTCTG |  |
| Ag_ rpS5_F | CCATGTCACGTCTCGTCACT | 104 |
| Ag_ rpS5_R | CGAAAACCATCCACACACAC |  |
