## Supplemental Figures for "Programmable CRISPR interference for gene silencing using Cas13a in mosquitoes"

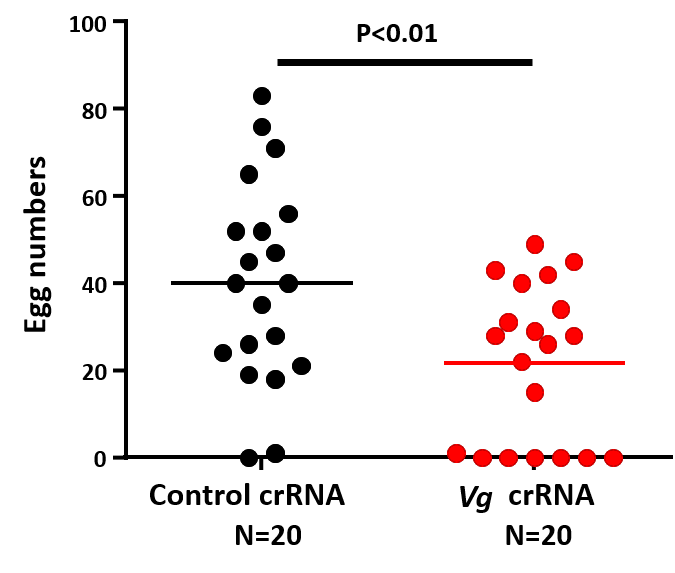

**Figure S1**. The Cas13a/*Vg* crRNA treatment significantly reduced egg production in *An. gambiae.* Mann-Whitney test, P<0.01.

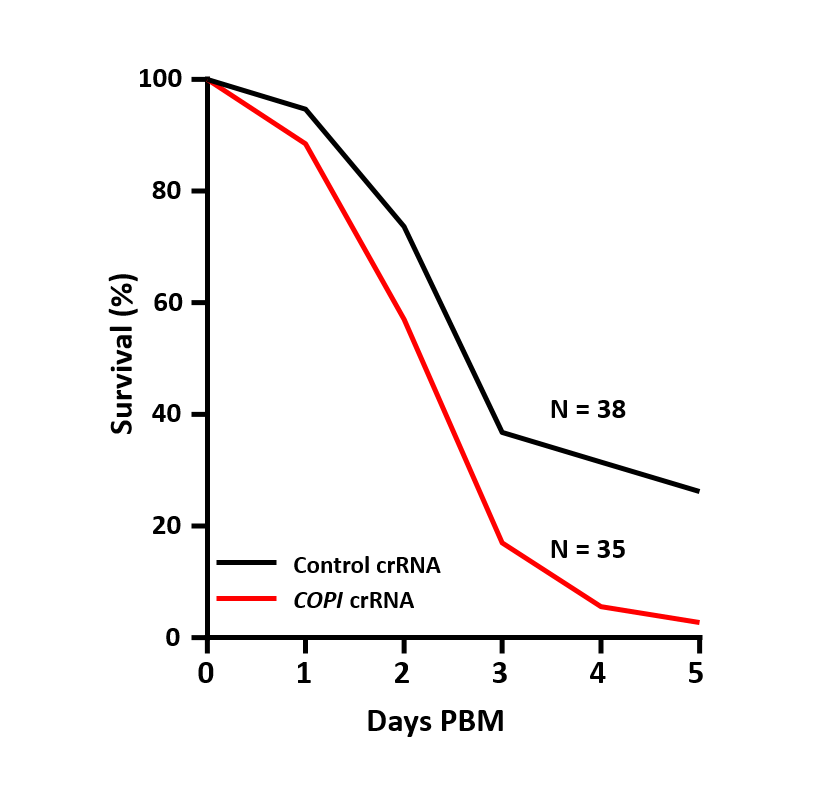

**Figure S2.** The Cas13a/*COPI-*crRNA treatment resulted in a significant higher mortality post a blood meal. Mantel Cox test, P<0.01.
